# Chemotaxis allows bacteria to overcome host-generated reactive oxygen species that constrain gland colonization

**DOI:** 10.1101/216515

**Authors:** Kieran D. Collins, Shuai Hu, Helmut Grasberger, John Y. Kao, Karen M. Ottemann

## Abstract

The epithelial layer of the gastrointestinal tract contains invaginations, called glands or crypts, which are colonized by symbiotic and pathogenic microorganisms and may function as designated niches for certain species. Factors that control gland colonization are poorly understood, but bacterial chemotaxis aids occupation of these sites. We report here that a *Helicobacter pylori* cytoplasmic chemoreceptor, TlpD, is required for gland colonization in the stomach. *tlpD* mutants demonstrate gland colonization defects characterized by a reduction in the percent of glands colonized, but not in number of bacteria per gland. Consistent with TlpD’s reported role in reactive oxygen species (ROS) avoidance, *tlpD* mutants showed hallmarks of exposure to large amounts of ROS. To assess the role of host-generated ROS in TlpD-dependent gland colonization, we utilized mice that lack either the ability to generate epithelial hydrogen peroxide or immune cell superoxide. *tlpD* gland colonization defects were rescued to wild-type *H. pylori* levels in both of these mutants. These results suggest that multiple types of innate immune generated ROS production limit gland colonization and that bacteria have evolved specific mechanisms to migrate through this gauntlet to establish in the glands.

**Classification:** Biological sciences; microbiology

**Significance statement:** Microbial colonization of the gastrointestinal tract occurs at distinct sites within the tissue including glandular structures found in the stomach and intestine. Multiple lines of evidence suggest that glands supply niches that promote chronic microbial colonization, a process that is critical for symbiotic and pathogenic bacteria to maintain themselves. In this report, we show that host-produced reactive oxygen species (ROS) constrain gland colonization by the gastric pathogen *Helicobacter pylori.* A bacterial cytoplasmic chemoreceptor, TlpD, allows *H. pylori* to avoid ROS and enhances *H. pylori*’s ability to colonize a broad swath of glands. We propose that hosts limit gland access and spread by producing ROS, and bacteria counter with chemotactic responses that allow navigation through this gauntlet.

## Introduction

The epithelium of the gastrointestinal (GI) tract contains invaginations, called glands in the stomach and crypts in the intestine, which are thought to serve as a niche for particular microbes and in turn, promote chronic colonization by specific microbial species. Our knowledge of the factors that control the colonization of these structures is incomplete. Host factors that have been implicated in controlling gland colonization include the production of mucus (1), the production of antimicrobial peptides (2), and the presence of resident immune cells in the lamina propria (3). Gland colonization, therefore, requires microbes to bypass these defensive strategies.

Bacteria too appear to have special adaptations to the gland niche. These include the ability to use certain carbohydrates (4) and perform chemotaxis (5–7). The chronically-colonizing gastric pathogen *Helicobacter pylori* is one such microbe that requires chemotaxis for gland colonization (5–7). Chemotaxis permits bacteria to sample their environment via chemoreceptors that use ligand-binding signals to alter the autophosphorylation of a complexed histidine kinase CheA. Ultimately, this pathway alters flagellar motility to allow bacteria to follow or repel themselves from gradients of specific signals (8). *H. pylori* expresses four chemoreceptors, three of which (TlpA, TlpB, and TlpC) are embedded the inner membrane, and one that is fully cytoplasmic (TlpD). The relevance of individual chemoreceptors on overall gastric colonization has been gauged previously by the level of colonization defect that a particular mutant displays. Among individual *H. pylori* chemoreceptor mutants, *tlpD* mutants display the most severe colonization attenuation in two animal models of infection (9, 10). The exact nature of the *tlpD* mutant colonization deficit, however, has remained unclear, as has the role of specific signals and chemoreceptors in gland colonization.

TlpD has been linked to a chemotactic response to multiple stress-related signals including electron transport chain inhibitors (11), acid (7), and reactive oxygen species (ROS) including hydrogen peroxide (H_2_O_2_) or the superoxide generators metronidazole and paraquat (12). One hypothesis is that these signals are connected because they all affect oxidative stress experienced in the cytoplasm (12). Gastric *Helicobacter* are known to encounter host-generated ROS derived from both epithelial and immune cells during infection and must cope with this stress to successfully colonize (13, 14).

ROS are produced by both gastric epithelial cells and innate immune cells, and include hydrogen peroxide (H_2_O_2_), superoxide (O_2^−^_) and hypochlorous acid (HOCl) (13). To counter these stresses, *H. pylori* possesses a suite of ROS detoxification systems including catalase, superoxide dismutase, and peroxiredoxins (15). *H. pylori* mutants lacking these systems are sensitive to ROS and are also attenuated in the host (15). ROS production limits colonization at epithelial surfaces in the stomach and intestine (14, 16); in agreement with this idea, mouse mutants that lack the epithelial DUOX enzyme produce less H_2_O_2_ and allow elevated colonization by a relative of *H. pylori*, *Helicobacter felis* (14). ROS production may serve to drive microbes away from the epithelial surface, as microbial adherence to intestinal epithelial cells promotes H_2_O_2_ production and hosts respond to *H. pylori* infection with elevated ROS (15, 16). However, it is not clear how ROS affects colonization within the glands.

To define the contribution of TlpD in gastric colonization, we first determined its effect on bacterial distribution in the stomach. We found that *tlpD* mutants showed specific deficits in colonizing a broad swathe of gastric glands, and displayed hallmarks of exposure to elevated ROS. This result raised the possibility that gland colonization defects could be due to an inability of *tlpD* mutants to successfully navigate in response to ROS, an idea that was further supported by the observation that tlpD mutants achieved normal numbers per gland in the glands they colonized. To assess whether host-generated ROS impacted *H. pylori* colonization, we compared the colonization and distribution of wild type (WT), *tlpD* and nonchemotactic *cheY* mutants in mice deficient in either epithelial dual oxidases (*Duoxa-/-*) or phagocytic NOX2 NADPH oxidase (*Cybb-/-)*. Infection of either *Duoxa-/-*or *Cybb-/-* mutant mice rescued the gland colonization defects of *tlpD* mutants noted in WT hosts. Our results suggest that ROS production impacts *H. pylori* gland transit, and that TlpD-mediated chemotactic responses are needed to thread this restricted gland access.

## Results

### tlpD mutants have minor colonization defects but achieve normal per gland loads

To begin our analysis of TlpD’s role in colonization, we orally infected WT C57BL6 mice with WT, *tlpD,* or *cheY* mutant variants of *H. pylori* that all expressed GFP. *cheY* encodes the central chemotaxis signaling proteins, so mutants that lack it are fully non-chemotactic, while mutants that lack *tlpD* lose only responses sensed by that receptor and thus are partially chemotactic. After two weeks of infection, the total colonization levels in tissue of the stomach corpus and antrum were determined. *tlpD* mutants showed colonization defects in the antrum and corpus of WT mice (Fig. 1A), similar to that previously reported (9). These results suggested that *tlpD* GFP+ *H. pylori* behaved similarly to *tlpD* infections lacking GFP described previously (9), and encouraged the analysis of gland colonization by the mutant.

**Figure 1.**
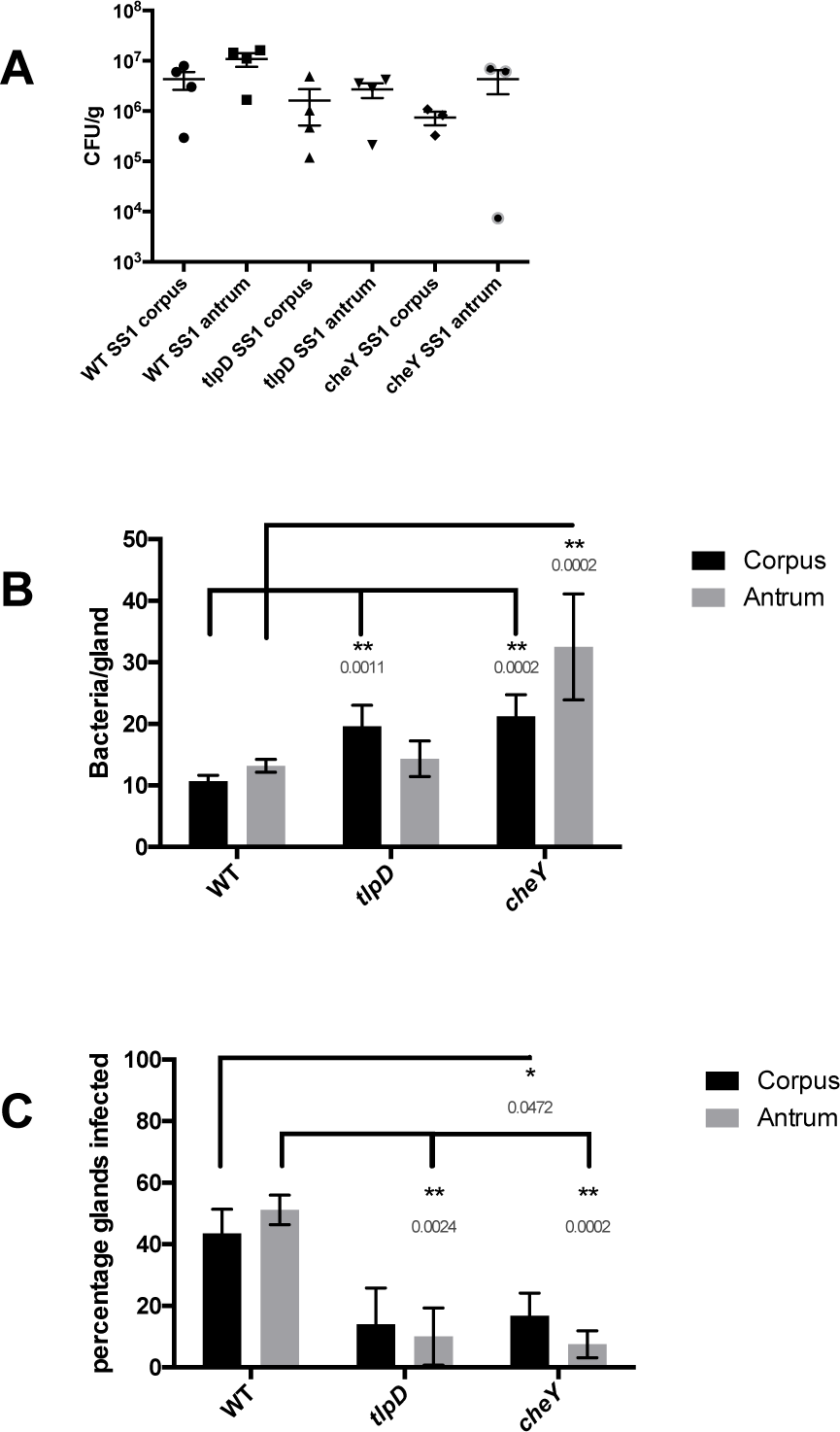
*tlpD* mutants have deficits in gland occupancy in WT mice but not colonization of total tissue or individual glands. Comparison of colonization of WT mice by *H. pylori* GFP+ SS1 WT, *tlpD*, and *cheY*. Mice were orally infected, and stomachs were collected and analyzed for tissue and gland colonization after 2 weeks of infection. (A) CFU/gram for corpus or antrum regions. *H. pylori* GFP+ SS1 WT (n = 4), *tlpD* (n = 4) and *cheY* (n = 3). (B) Gland loads in the isolated corpus and antral glands. These numbers are the average number of bacteria counted per gland, excluding uninfected glands. Infected gland numbers are: WT corpus (436 glands from 6 mice), WT antrum (508 from 6 mice), *tlpD* corpus (67 glands from five mice), *tlpD* antrum (48 glands from five mice), *cheY* corpus (58 glands from four mice), *cheY* antrum (24 glands from four mice). (C) Gland occupancy in the isolated corpus and antral glands, representing the percentage of glands infected with the indicated *H. pylori* strain. Error bars represent standard error of the mean (SEM) for all panels. Numbers of mice infected are the same as described for gland loads. Statistical differences are indicated by * (*P <* 0.05) and ** (*P <* 0.01) as analyzed by Student T-test.

We next sought to examine TlpD’s role in gland colonization. To monitor gland colonization, we employed the bacterial localization in isolated glands (BLIG) approach in which gastric glands are isolated from the infected corpus or antrum tissue, epithelial cells labeled with Hoechst DNA stain, and glands examined for the presence of GFP+ *H. pylori* by fluorescent microscopy (5). Bacteria within glands were counted manually, and two parameters of gland colonization were compared between *H. pylori* strains. The first parameter was gland bacterial load, the number of bacteria per infected gland. Our calculation of gland bacterial load excludes non-infected glands. The second parameter was gland occupancy, the percent of glands infected.

In WT mice, WT *H. pylori*-colonized the glands of both the corpus and the antrum to similar levels, averaging 10 bacteria/infected gland as reported previously (Fig. 1B) (5). Loss of TlpD did not affect gland load in the antrum but caused a ~1.8-fold increase in gland load in the corpus compared with WT (Fig. 1B). Full loss of chemotaxis (*cheY* mutants) also resulted in elevated gland loads of 2-to 3-fold in both the corpus and the antrum relative to WT *H. pylori* (Fig. 1B). These results suggest that chemotactic defects did not impair growth within glands, and if anything, resulted in elevations in bacterial gland load.

### The loss of *tlpD* or chemotaxis results in a reduction in gland occupancy throughout the stomach in WT hosts

Because *tlpD* and *cheY* mutants appeared to have altered gland phenotypes, we next analyzed gland occupancy to determine the percentage of glands infected by *H. pylori*. This frequency likely reflects both the initial population of glands infected by *H. pylori* as well as the ability to spread and colonize new glands. In WT mice, WT *H. pylori* colonized 40-50% of corpus and antral glands by two weeks of infection, and were found in similar proportions in both regions (Fig. 1C). *tlpD* mutants showed an ~3-fold reduced occupancy in both the corpus and antrum relative to WT *H. pylori* (Fig. 1C). *cheY* mutant gland occupancy was also decreased relative to WT *H. pylori*, with significant reductions in both the corpus and the antrum (Fig. 1C). These results suggest that chemotaxis generally and TlpD specifically is required for *H. pylori* to occupy new glands.

### tlpD mutants show hallmarks of elevated ROS exposure relative to WT *H. pylori*

We reported recently that TlpD mediates chemotactic repellent responses to multiple ROS (12). Combining this information with our data above suggested that *tlpD* mutant gland colonization defects could be due to an inability of these mutants to sense and repel themselves away appropriately from ROS. We therefore asked whether *tlpD* mutants experienced differential oxidative stress *in vivo*. For this approach, we used quantitative real-time PCR of mRNA isolated from infected mouse tissue. We examined the expression of the catalase gene (*katA*) mRNA by *H. pylori* strains, whose expression has been shown to be sensitive to several oxidative stresses (17, 18). We determined that this gene was modestly upregulated *in vitro* in our strains following exposure to 1 mM H_2_O_2_ for twenty minutes (Fig. 2A). This result suggested that *katA* mRNA could serve as a reasonable proxy for H_2_O_2_ exposure *in vivo*. We next assessed whether the expression of *katA* mRNA differed between WT, *tlpD,* or *cheY H. pylori* during infection of WT mice. *tlpD* mutants expressed significantly more *katA* mRNA than WT *H. pylori* in the antrum, and modestly more in the corpus (Fig. 2B). These results suggest that *tlpD* mutants experience elevated oxidative stress during infection. Conversely, *cheY* mutants did not express elevated catalase mRNA (Fig 2B). This outcome suggests that the loss of TlpD specifically leads *H. pylori* to be exposed to conditions that are different than those encountered by WT, consistent with high exposure to oxidative stress.

**Figure 2.**
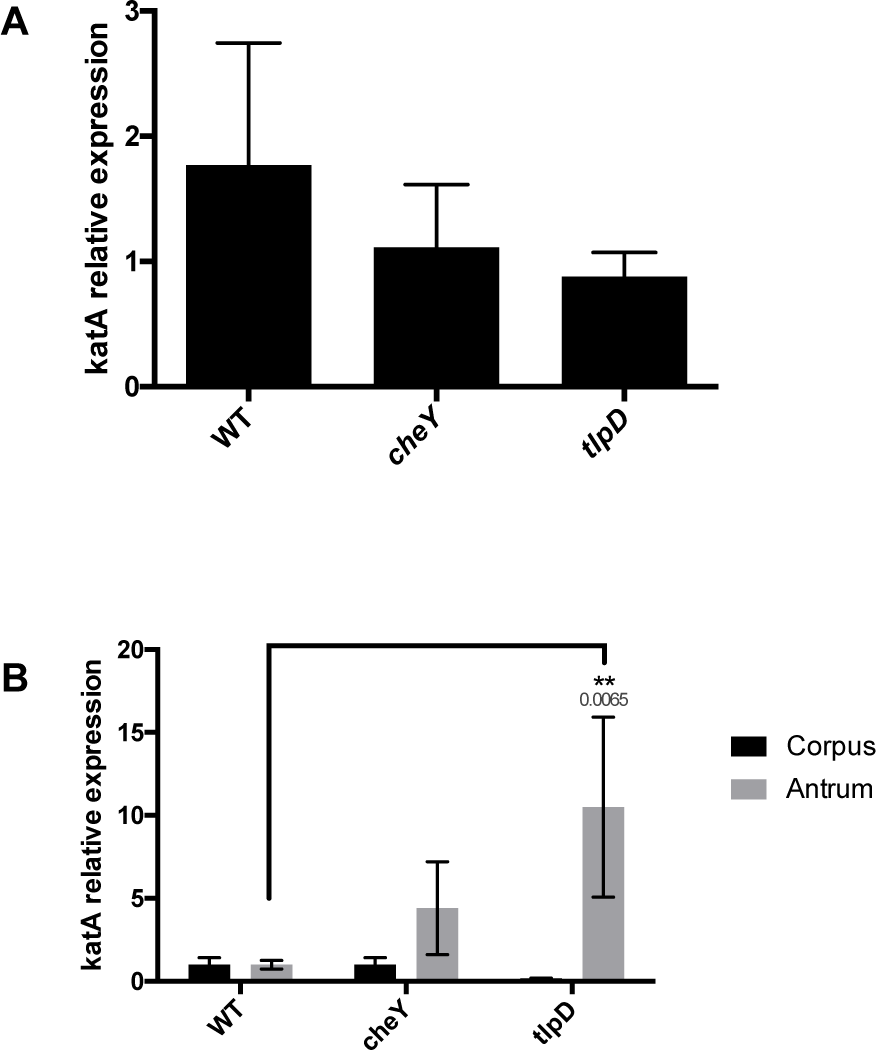
*tlpD* mutants show evidence of ROS exposure *in vivo*. Comparison of catalase mRNA expression *in vitro* and *in vivo* between *H. pylori* strains.(A) Mean +/-SEM of fold change increases in *katA* mRNA of *H. pylori* strains exposed to 1 mM H_2_O_2_ for twenty minutes, normalized to *gapB*. (B) Comparison of mean +/-SEM of *katA* expression by *H. pylori* strains in three WT mice, normalized to *gapB*. Statistical differences are indicated by * (*P <* 0.05) and ** (*P <* 0.01) as analyzed by Student T-test, with actual p values indicated above the bar. *gapB* expression was insensitive to H_2_O_2_ exposure based on comparison to 16S rRNA.

### Gland colonization defects of *tlpD* are rescued in hosts deficient in H_2_O_2_ production by gastric epithelial cells

The results presented above suggest that TlpD helps to mitigate exposure of *H. pylori* to oxidative stress in the mouse. In order to follow up on oxidative stress and its role in TlpD-mediated colonization, we next infected two mutant mouse hosts that were deficient in the production of H_2_O_2_ and O_2_-production. The first of these lacks the dual oxidase (Duox) heterodimeric enzyme complex by virtue of loss of the *Duoxa*-encoded subunit (14). Duox is expressed by gastric epithelial cells and generates extracellular H_2_O_2_ that may serve to limit physical interactions between microbes and the epithelial surface (16). Duox has been implicated in limiting the colonization of a related *Helicobacter* species in the stomachs of mice (14).

To examine whether Duox impacted *H. pylori* colonization, *Duoxa-/-* mice were infected as done with WT mice for two weeks, at which point the mice were sacrificed and colonization of WT, *cheY,* and *tlpD* GFP+ *H. pylori* was compared. All *H. pylori* strains colonized the *Duoxa -/-* mutants to levels that were a bit elevated but not significantly different from those in WT mouse hosts (Fig. 3A). Gland loads were also generally similar between WT and *Duoxa-/-* glands, across WT and *tlpD* mutant *H. pylori* in both locations, and *cheY* mutants in the corpus (Fig. 3B). There was a modest increase in gland load in the antrum of the *tlpD* mutant and a very large decrease in loads of the *cheY* mutant, suggesting the effect of Duox was greatest in the antrum.

**Figure 3.**
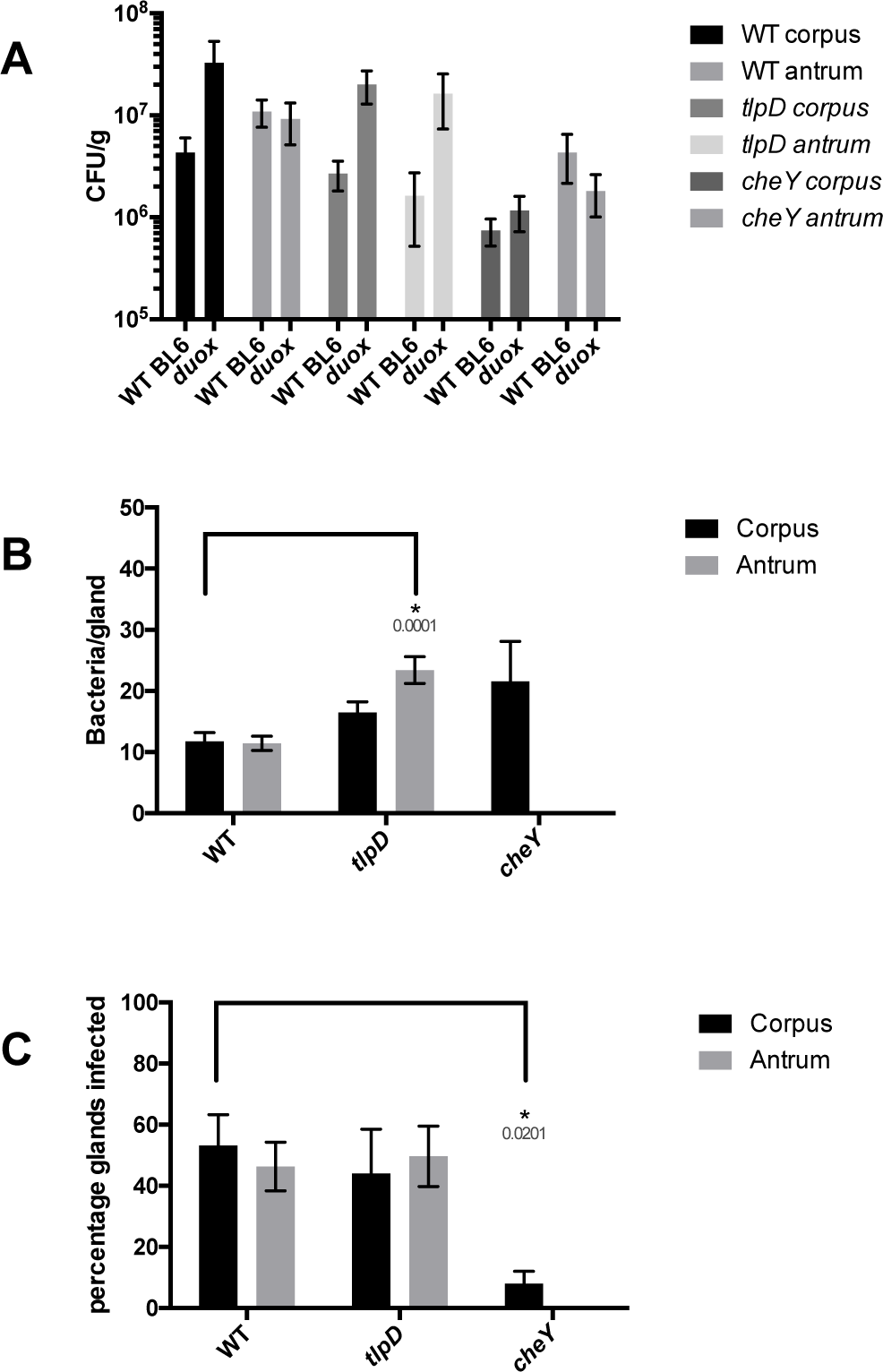
Loss of epithelial H_2_O_2_ rescues *tlpD* mutant gland defects. Colonization of *Duoxa-/-* mice by WT, *tlpD,* and *cheY* GFP+ *H. pylori* SS1 strains at two weeks post-infection. Mice were orally infected, and stomachs were collected and analyzed for tissue and gland colonization. (A) CFU/gram at two weeks post-infection for corpus or antrum regions with WT (n = 4), *tlpD* (n = 5) and *cheY* (n = 5) GFP+ *H. pylori* SS1. Data for WT mice are the same as in Fig. 1, and are reshown here for comparison. (B) Gland loads in the isolated corpus and antral glands, representing the average number of bacteria counted per gland, excluding uninfected glands. Infected gland numbers are: WT corpus (313 glands from six mice), WT antrum (472 glands from 6 mice), *tlpD* corpus (132 from six mice), *tlpD* antrum (149 glands from three mice), *cheY* corpus (24 glands from three mice). (C) Gland occupancy in the isolated corpus and antral glands, representing the percentage of glands infected with the indicated *H. pylori* strain. Error bars represent SEM for all panels. Numbers of mice infected are the same as described for gland loads. Statistical differences are indicated by * (*P <* 0.05) and ** (*P <*0.01) as analyzed by Student T-test.

We next assessed how the loss of *Duoxa-/-* would alter gland occupancy. WT *H. pylori* gland occupancy was seemingly unaffected by the loss of *Duoxa-/-*, as ~50% of glands were infected in this background as well as in WT mice (Fig. 3C). Interestingly, the *tlpD* mutant showed an increase in gland occupancy compared to its levels in a WT mouse, moving from <15% occupied to over 40% (Fig. 3C). Indeed, the *tlpD* mutant achieved gland occupancy levels in the corpus and antrum that were not different from WT *H. pylori* (Fig. 3C). In contrast, the *cheY* mutant was not rescued, suggesting the loss of *Duoxa-/-* rescue is specific to signals sensed by TlpD and requires chemotaxis. This apparent rescue in *tlpD* gland occupancy suggests that the loss of H_2_O_2_ production by gastric epithelial cells allows for *tlpD* mutants to move more readily into new gastric glands in both the corpus and the antrum.

### Gland colonization defects of *tlpD* are rescued in hosts deficient in O_2_- production by phagocytes

We next assessed the contribution of phagocyte ROS production to *H. pylori* gland colonization. Phagocyte ROS production was assessed in *Cybb-/-* mice that lack the catalytic subunit of phagocyte oxidase (Phox). *Cybb-/-* mice were infected as above and the same colonization parameters were compared between WT, *cheY,* and *tlpD* GFP+ *H. pylori*.

The overall colonization of the corpus and antrum was seemingly unaffected by loss of *Cybb* for all three *H. pylori* strains, showing no significant differences from WT mouse infections (Fig. 4A). Gland loads, on the other hand, were affected in *Cybb-/-* hosts. Both WT and *tlpD H. pylori* showed elevated gland loads in the corpus and the antrum relative to WT BL6 infections, achieving 20-30 bacteria/gland in both regions. *cheY* mutants did not follow this trend in *Cybb-/-* hosts and instead showed reduced gland loads in the corpus and the antrum relative to WT BL6 infections (Fig 4B). This outcome suggests that superoxide may limit *H. pylori* numbers in a chemotaxis-dependent way. Lastly, we compared gland occupancy in *Cybb-/-* hosts. Strikingly, the *tlpD* mutant gland occupancy in both the corpus and antrum climbed to levels that were not different from WT *H. pylori*. This finding suggests that, similarly to *Duoxa-/-* infections, gland occupancy defects of *tlpD* were rescued by loss of host ROS. As seen with *Duoxa-/-* infections, *cheY* gland occupancy did not appear to benefit from the loss of *Cybb-/-* (Fig 4C). These results suggest that the loss of superoxide production by phagocytes rescues gland colonization defects of *tlpD H. pylori*, as was observed in *Duoxa-/-* hosts. Chemotaxis appears necessary for this rescue, as *cheY* mutants showed similar gland colonization values observed in WT mice. Taken together these results suggest that host-generated ROS serves as a barrier for gland colonization by *H. pylori* that the bacteria overcome via TlpD-mediated chemotactic responses. Furthermore, *tlpD* colonization defects can be attributed to low gland occupancy in the corpus and the antrum, which can be rescued to WT levels by disrupting host ROS production.

**Figure 4.**
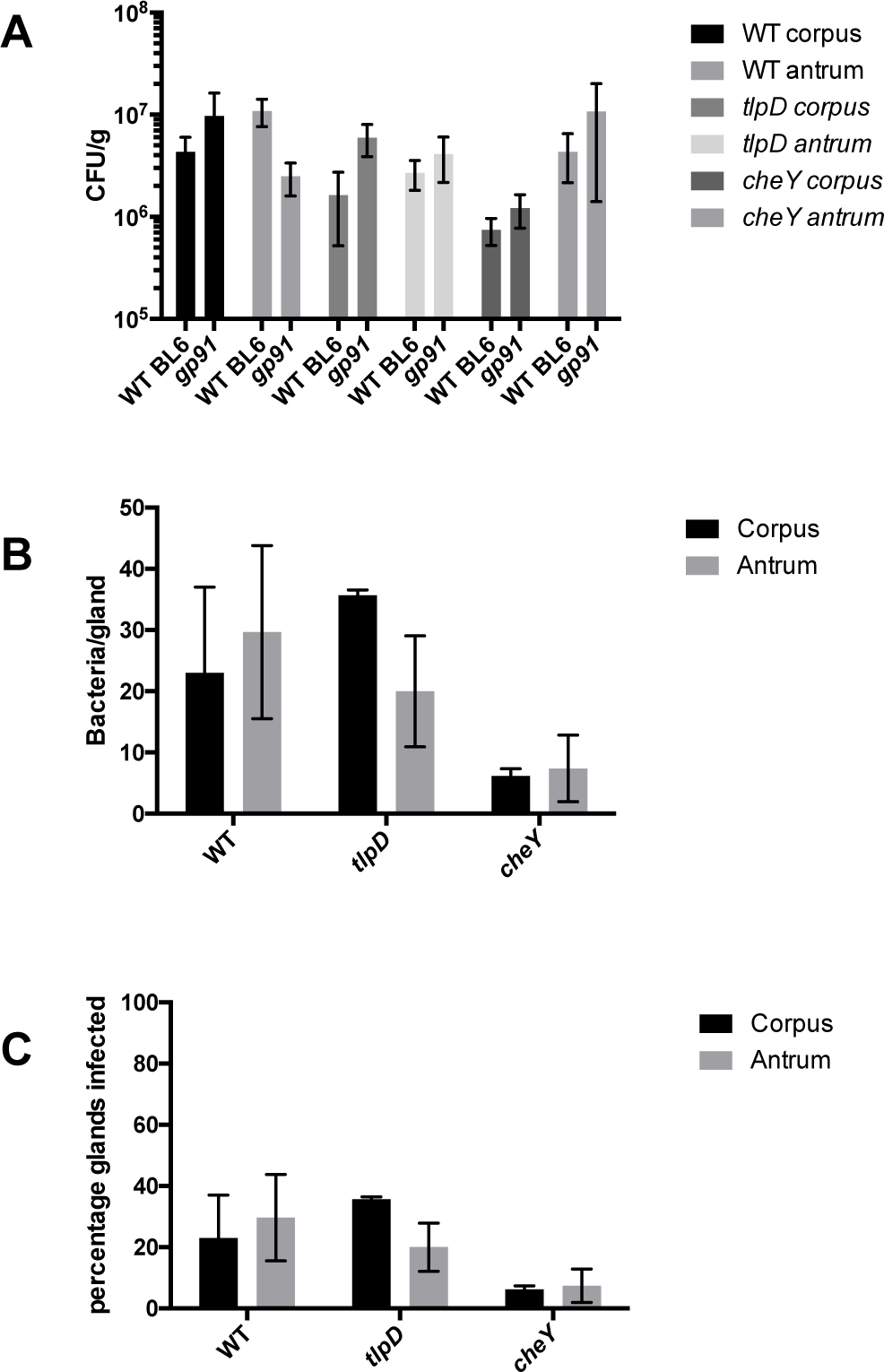
Loss of immune superoxide rescues *tlpD* mutant gland defects. Colonization of *Cybb-/-* mice by WT, *tlpD*, and *cheY* GFP+ *H. pylori* SS1 strains at two weeks post-infection. Mice were orally infected, and stomachs were collected and analyzed for tissue and gland colonization. (A) CFU/gram at two weeks post-infection for corpus or antrum regions using WT (n = 6), *tlpD* (n = 14) and *cheY* (n = 6) GFP+ *H. pylori* SS1 strains. Data for WT mice are the same as in Fig. 1, and are reshown here for comparison. (B) Gland loads in the isolated corpus and antral glands, representing the average number of bacteria counted per gland, excluding uninfected glands.Infected gland numbers are: WT corpus (69 glands from three mice), WT antrum (89 glands from 3 mice), *tlpD* corpus (107 glands from three mice), *tlpD* antrum (60 glands from three mice), *cheY* corpus (31 glands from three mice), *cheY* antrum (37 glands from three mice). (C) Gland occupancy in the isolated corpus and antral glands, representing the percentage of glands infected with the indicated *H. pylori* strain.Error bars represent SEM for all panels. Numbers of mice infected are the same as described for gland loads. Statistical differences are indicated by * (*P <* 0.05) and ** (*P <* 0.01) as analyzed by Student T-test.

## Discussion

We report here the *H. pylori* requires repellent ROS chemotaxis to be able to successfully colonize glands. Factors that control gland colonization throughout the GI tract are poorly understood, although it appears that an interplay exists between host and microbe to regulate gland access. Host factors known to limit gland colonization include mucus production (1), oxygen gradients emanating from the epithelial surface (19), and antimicrobial peptide production (2). Microbial adaptations that have been reported to aid gland colonization include chemotaxis (5–7), sugar transport systems (20), and the ability to dampen host immune responses (21). Therefore it seems reasonable to posit that glands represent a desired but protected niche for some microbes in the GI tract. Our work demonstrates that ROS limits gland access, and chemotaxis helps overcome this barrier.

Host ROS generation has been implicated in limiting microbial adhesion in the intestine, and *Duoxa-/-* mice showed elevated mucosal penetrance by a subset of the microbiota (16). Our data suggest that host ROS plays an important role in restricting gland access in the stomach, and that bacteria can use chemotaxis to overcome this barrier. Gland colonization defects observed for *tlpD* mutants in WT hosts were effectively rescued in hosts with ROS production defects. Low gland occupancy could be due to very low initial colonization, or to low gland-gland spread, as both of these processes benefit from chemotaxis (5). Our results lend support to a prior report suggesting that TlpD mediates chemotactic repellent responses to ROS treatments *in vitro* (12), and defines the nature of colonization defects of *tlpD* mutants which have been described in the past (9, 10).

Our work additionally suggests that chemotaxis is not required for growth once bacteria are in glands, because we observed here that non-chemotactic and *tlpD* mutants obtained high numbers/gland in WT mice. These results are match those in a prior report, which presented the average number of bacteria/gland in a manner that included uninfected glands in that calculation (5). Excluding uninfected glands from data reported in Keilberg *et al*. concerning gland loads for WT and *cheY H. pylori* would produce similar values as those described in this report (5). It is not yet known what sets gland load, but it has been observed that this number varies over the course of a mouse infection, climbing to an average of ~15-25 bacteria/gland within the first month, and then dropping to less than 5 by six months of infection (5). Our results show that chemotaxis can affect these within-gland levels, somewhat surprisingly playing a role to limit bacterial numbers. Our data suggest the possibility that chemotaxis plays a critical role in gland exit, such that without chemotaxis or TlpD specifically, bacterial numbers rise in the glands but bacteria cannot effectively leave. This phenotype in turn creates poor gland occupancy.

Previous work showed that TlpD drives chemotactic repellent responses i*n vitro* (7, 11, 12), and our data is consistent with the idea that it also mediates chemotactic repellent responses in the host. Specifically, we found that *tlpD* mutants display hallmarks of high ROS exposure, in agreement with the idea that these mutants cannot avoid ROS. One role for this response *in vivo* comes from the observation that hosts upregulate defensive ROS production upon *H. pylori* infection (15). Thus, *H. pylori* may experience a delay from initially colonizing glands to experiencing stress imparted by the host. Therefore a repellent response mediated by TlpD could limit the detrimental effect of these stresses.

In conclusion, we have described host ROS generation as an additional host limitation on gland colonization in the stomach that is overcome by chemotaxis. We implicate the *H. pylori* cytoplasmic chemoreceptor TlpD in ROS-dependent gland colonization effects in the host and show that colonization defects noted for a *tlpD* mutant in WT hosts is relieved in ROS-production deficient hosts. TlpD appears to be involved in the dispersal of *H. pylori* between glands in a ROS-dependent fashion.

## Acknowledgements

The authors thank Christina Yang for contributing references and discussion about factors that affect gland colonization. The described project was supported by National Institutes of Health National Institute of Allergy and Infectious Disease (NIAID) grant R21AI117345 (to K.M.O.) and RO1DK087708-01 (to J.Y.K), as well as UCSC Committee on Research funds (to K.M.O). The funders had no role in study design, data collection and interpretation, or the decision to submit the work for publication.

## Methods

### Bacterial strains and culture conditions

WT and *cheY* SS1 GFP+ *H. pylori* described previously were employed for mouse infections (5). *H. pylori* was cultured on either Columbia horse blood agar (CHBA), or brucella broth with 10% fetal bovine serum (FBS; Life Technologies) (BB10). CHBA consisted of Columbia agar (BD) with 5% defibrinated horse blood (Hemostat Labs, Davis, CA), 50 µg/ml cycloheximide, 10 µg/ml vancomycin, 5 µg/ml cefsulodin, 2.5 U/ml polymyxin B, and 0.2% (wt/vol) β-cyclodextrin. All chemicals were from Thermo Fisher or Gold Biotech. Cultures were incubated at 37°C under 5 to 7% O_2_, 10% CO_2_, and balance N_2_.

### Creation of *tlpD* GFP+ *H. pylori* mutants

*tlpD* GFP+ *H. pylori* strain SS1 (KO1614) was constructed by transformation of ∆*tlpD::cat* SS1 (KO914) (12, 22) with the plasmid pTM115 (5, 23) isolated from *H. pylori* strain SS1, and selected on CHBA plates containing 15 ug/ml kanamycin (5, 9).

### Animal infections and *H. pylori* colonization calculations

The University of California, Santa Cruz Institutional Animal Care and Use Committee approved all animal protocols and experiments. *Cybb-/-* targeted homozygous null mice in a B6.129S background were obtained from Jackson Laboratory (JAX stock #002365, Bar Harbor, ME)(24); *Duoxa*-/-mice lacking functional dual oxidase enzymes by virtue of loss of the duoxa1-duoxa2 maturation subunits (25) were obtained as heterozygotes on the B6 background from the University of Michigan. All mice were obtained as breeding pairs, and bred at UC Santa Cruz. *Duoxa*-/-mice were generated, screened, and maintained as previously described (14). In brief, *Duoxa* genotyping was performed by isolating genomic DNA from tail tissue with the Qiagen DNeasy Blood & Tissue Kit., followed by PCR with a common primer (DA-WT/KO), a WT allele-specific primer (DA-WT-R), and a knockout allele-specific primer (DA-KO-R) (14). Genotypes were judged by the presence of the WT allele as a 381-basepair fragment, and the knockout allele as a size of 568 basepair fragment (25).

Six to eight-week-old mice (male and female) were infected intraorally by allowing the animals to drink a 50 microliter suspension from a pipette tip containing *H*. *pylori* grown to mid-exponential phase and concentrated to an optical density at 600 nm of 3.0 (~ 5x107/50 µl) in BB10 medium, as done previously (6). At the end of an infection period, mice were sacrificed by CO_2_ narcosis. The stomach was removed, opened along the lesser curvature and washed in phosphate-buffered saline (PBS) to remove food. The corpus and antrum were divided based on tissue coloration, cut into pieces that were then processed to analyze total bacterial colonization, gland isolation, or for RNA extraction. For total bacterial colonization, corpus and antral tissue was weighed, homogenized with the Bullet Blender (Next Advance) with 1.0-mm zirconium silicate beads, and then plated to determine the number of colony forming units (CFU) per gram of stomach tissue on CHBA with the addition of 20µg/ml bacitracin, 10 µg/ml nalidixic acid, and 15 µg/ml kanamycin.

### Gland isolation and microscopy

Glands were isolated by incubating dissected gastric tissue in Dulbecco’s phosphate-buffered saline (DPBS) (Millipore) plus 5 mM EDTA at 4°C for 2 hours with agitation, as described previously (5, 26). The tissue was subsequently transferred to DPBS containing 1% sucrose and 1.5% sorbitol and shaken for thirty seconds. Glands were labeled with 10 μg/ml Hoechst DNA stain (Life Technologies). Glands were kept on ice and examined as soon as possible. Ten microliters of shaken tissue were placed on glass slides and visualized with a Nikon Eclipse E600 microscope with fluorescence filters for 4’,6’-diamidino-2-phenylindole (DAPI), GFP, and RFP. For each time point and infection, 100 glands each were imaged for the corpus and antrum, and the number of *H. pylori* cells inside the gland was counted manually for each gland. Gland load levels were calculated by averaging the number of bacteria observed in colonized glands per mouse and *H. pylori* strain. Gland occupancy was calculated as the frequency of glands occupied per mouse host and averaged over at least three mice. Gland colonization comparisons were made for at least three mice per genotype and *H. pylori* strain.

### RNA isolation and qPCR

Gastric tissue was flash frozen in liquid nitrogen, homogenized in TRIzol (Invitrogen) and RNA was isolated following the TRIzol RNA isolation protocol (GIBCO). DNA was removed by following the TURBO DNA-free kit protocol (Life technologies). cDNA was produced with the High-Capacity cDNA Reverse Transcription Kit (Life technologies) using random primers. qPCR was performed using the SensiFAST SYBR No-ROX kit (Bioline) using the primers listed below. Primer efficiency was calculated by amplifying serial dilutions of WT *H. pylori* genomic DNA, plotting the Ct values obtained per dilution and calculating the slope. Efficiencies were derived from the slope with the equation Efficiency = −1+10^(−1/slope)^ (27). Relative fold changes were calculated using the ΔΔCt method with Pfaffl correction for PCR amplification efficiency, using *16S* and *gapB* as reference genes with primers listed 5’−3’ below (27). 16S forward: GGAGGATGAAGGTTTTAGGATTG; 16S reverse: TCGTTTAGGGCGTGGACT*; katA* forward: AGAGGTTTTGCGATGAAGT*; katA* reverse: CGTTTTTGAGTGTGGATGAA*; gapB* forward: GCCTCTTGCACGACTAACGC*; gapB* reverse: CTTTGCTCACGCCGGTGCTT.

### *In vitro* treatment of *H. pylori* with H_2_O_2_

Overnight cultures of *H. pylori* strains were adjusted to OD_600_ = 0.2, split into two cultures with one receiving treatment with 1 mM H_2_O_2_ for twenty minutes. RNA isolation and qPCR protocols were identical to that described above.

## References

1. Millet YA, et al. (2014) Insights into Vibrio cholerae Intestinal Colonization from Monitoring Fluorescently Labeled Bacteria. PLoS Pathog 10(10):e1004405.

2. Petnicki-Ocwieja T, et al. (2009) Nod2 is required for the regulation of commensal microbiota in the intestine (National Acad Sciences), pp 15813–15818.

3. Macpherson AJ, Slack E, Geuking MB, McCoy KD (2009) The mucosal firewalls against commensal intestinal microbes. Semin Immunopathol 31(2):145–149.

4. Lee SM, et al. (2013) Bacterial colonization factors control specificity and stability of the gut microbiota. Nature 501(7467):426–429.

5. Keilberg D, Zavros Y, Shepherd B, Salama NR, Ottemann KM (2016) Spatial and Temporal Shifts in Bacterial Biogeography and Gland Occupation during the Development of a Chronic Infection. mBio 7(5). doi:10.1128/mBio.01705-16.

6. Howitt MR, et al. (2011) ChePep controls Helicobacter pylori Infection of the gastric glands and chemotaxis in the Epsilonproteobacteria. mBio 2(4). doi:10.1128/mBio.00098-11.

7. Huang JY, Goers Sweeney E, Guillemin K, Amieva MR (2017) Multiple Acid Sensors Control Helicobacter pylori Colonization of the Stomach. PLoS Pathog 13(1):e1006118.

8. Lertsethtakarn P, Ottemann KM, Hendrixson DR (2011) Motility and Chemotaxis in Campylobacter and Helicobacter. Annu Rev Microbiol 65(1):389–410.

9. Rolig AS, Shanks J, Carter JE, Ottemann KM (2012) Helicobacter pylori requires TlpD-Driven chemotaxis to proliferate in the antrum. Infection and Immunity 80(10):3713–3720.

10. Behrens W, et al. (2013) Role of Energy Sensor TlpD of Helicobacter pylori in Gerbil Colonization and Genome Analyses after Adaptation in the Gerbil. Infection and Immunity 81(10):3534–3551.

11. Schweinitzer T, et al. (2008) Functional characterization and mutagenesis of the proposed behavioral sensor TlpD of Helicobacter pylori. Journal of Bacteriology 190(9):3244–3255.

12. Collins KD, et al. (2016) The Helicobacter pylori CZB Cytoplasmic Chemoreceptor TlpD Forms an Autonomous Polar Chemotaxis Signaling Complex That Mediates a Tactic Response to Oxidative Stress. Journal of Bacteriology 198(11):1563–1575.

13. Handa O, Naito Y, Yoshikawa T (2010) Helicobacter pylori: a ROS-inducing bacterial species in the stomach. Inflamm Res 59(12):997–1003.

14. Grasberger H, Zaatari El M, Dang DT, Merchant JL (2013) Dual Oxidases Control Release of Hydrogen Peroxide by the Gastric Epithelium to Prevent Helicobacter felis Infection and Inflammation in Mice. Gastroenterology 145(5):1045–1054.

15. Flint A, Stintzi A, Saraiva LM (2016) Oxidative and nitrosative stress defences of Helicobacter and Campylobacter species that counteract mammalian immunity. FEMS Microbiology …. doi:10.1093/femsre/fuw025.

16. Grasberger H, et al. (2015) Increased Expression of DUOX2 Is an Epithelial Response to Mucosal Dysbiosis Required for Immune Homeostasis in Mouse Intestine. Gastroenterology 149(7):1849–1859.

17. Olczak AA, Wang G, Maier RJ (2009) Up-expression of NapA and other oxidative stress proteins is a compensatory response to loss of major Helicobacter pylori stress resistance factors. Free Radic Res 39(11):1173–1182.

18. Ernst FD, et al. (2005) Transcriptional profiling of Helicobacter pylori Fur-and iron-regulated gene expression. Microbiology (Reading, Engl) 151(Pt 2):533–546.

19. Pédron T, Nigro G, Sansonetti PJ (2016) From homeostasis to pathology: decrypting microbe–host symbiotic signals in the intestinal crypt. Phil Trans R Soc B 371(1707):20150500–4.

20. Lee SM, et al. (2013) Bacterial colonization factors control specificity and stability of the gut microbiota. Nature 501(7467):426–429.

21. Round JL, et al. (2011) The Toll-Like Receptor 2 Pathway Establishes Colonization by a Commensal of the Human Microbiota. Science 332(6):974–.

22. Williams SM, et al. (2007) Helicobacter pylori Chemotaxis Modulates Inflammation and Bacterium-Gastric Epithelium Interactions in Infected Mice. Infection and Immunity 75(8):3747–3757.

23. Amieva MR, et al. (2003) Disruption of the Epithelial Apical-Junctional Complex by Helicobacter pylori CagA. Science 300(5):1430–1434.

24. Pollock JD, et al. (1995) Mouse model of X–linked chronic granulomatous disease, an inherited defect in phagocyte superoxide production. Nature Genetics 9(2):ng0295–202–209.

25. Grasberger H, et al. (2012) Mice deficient in dual oxidase maturation factors are severely hypothyroid. Mol Endocrinol 26(3):481–492.

26. Mahe MM, et al. (2013) Establishment of Gastrointestinal Epithelial Organoids.

27. Pfaffl MW (2001) A new mathematical model for relative quantification in realtime RT-PCR. Nucleic Acids Research 29(9):45e–45.

